# ANALYSIS OF THE IMPACT OF FIVE METHANE–MITIGATING FEED ADDITIVES ON MILK PRODUCTION AND ASSOCIATED PARAMETERS ACROSS MULTIPLE COMMERCIAL DAIRY FARMS

**DOI:** 10.1101/2025.02.04.636425

**Authors:** Yaniv Altshuler, Tzruya Calvao Chebach, Shalom Cohen, Joao Gatica

## Abstract

The effect of methane–mitigating feed additives on dairy cows has been widely explored; however, confusing conclusions have been reached due to factors such as the inclusion of different doses and experimental conditions in which the additives are tested, or even a small sample size. We present the first extensive study assessing the effects of methane–mitigating feed additives on milk production across several commercial dairy farms. This study used a previously developed predictive AI–driven model based on microbiome samples; the model predicts farms where a significant reduction of methane emissions is expected due to the applied feed additives. Thus, in this study, each feed additive was supplied to a large number of farms, widely distributed across different climatic areas in Israel. The data analysis followed two simulated scenarios: (1) a naive approach, where feed additives are supplied indiscriminately, and (2) an optimized approach, where feed additives are supplied only to farms with a high likelihood of being positively impacted in terms of reduced enteric methane emissions (50% of the farms). The results show that each feed additive significantly increased milk production compared to the control groups. This increase in milk production was significantly higher in the optimized scenario. Other related parameters such as somatic cells were also improved. Our results suggest that the feed additives positively affect milk production, reaching a maximum expression when the AI–driven model is applied.

## INTRODUCTION

The dairy industry faces major global challenges, including advancing innovation in milk production, promoting sustainability to reduce environmental impact, and ensuring profitability and market competitiveness (Hassoun et al., 2023). Added to this, the increasing wealth of emerging economies has raised the demand for the supply of milk and meat (Silva et al., 2018). In this context, methane–mitigating feed additives offer an interesting response to those challenges, since according to different studies, these additives could reduce methane emissions, increase milk production, and provide defense against some diseases (Hart et al., 2019; Roque et al., 2019; Tseten et al., 2022). Despite extensive research in this field and some consensus on the effect of feed additives, their use often fails to deliver the expected results, especially when applied under standard dairy commercial conditions. These inconsistencies limit its implementation and diminish the overall impact of methane–mitigating feed additives (Roques et al., 2024). The heterogeneity of results can be explained by clear differences in the methodology used in such studies, including the low number of animals tested, laboratory conditions versus commercial conditions, the type and nature of the tested feed additives, and even the method of sampling and data recording (Calsamiglia et al., 2007).

In previous work oriented to mitigate enteric methane emissions in dairy farms, we developed a predictive AI–driven model that used microbiome data of several dairy farms in Israel to successfully predict the effect of methane–mitigating feed additives on methane emissions (Altshuler et al., 2023; Altshuler et al., 2025). This study proved the model’s accuracy but raised new questions such as the effect of the feed additives on milk production.

Thus, this study aims to compare the effects of five different methane-mitigating feed additives on milk production and related parameters, such as fat and protein content, lactose levels, and somatic cell count, under real commercial conditions across several farms. All the data used in this study was obtained by an automatized system (NOA) that monitors and registers the commercial milk collection. In addition, the system registers several parameters related to the milk quality, such as fat and protein content, lactose, and somatic cells, among others.

Each feed additive was tested on several commercial dairy farms across Israel. The selected commercial feed additives included four essential oil–based products and one monensin–based product.

## MATERIALS AND METHODS

### General study design

In this study, five feed additives were tested across a large number of farms, ranging from 13 to 21 farms for each additive, with each farm housing between 400 to 900 cows. To avoid seasonal effects, the participating farms were widely distributed across different climatic regions in Israel. On each site, the farm staff randomly selected an average of 15 cows for the treatment group, identifying the selected cows by their ear tag, and received a specific feed additive according to the manufacturer’s recommendations while remaining housed in the same barn as the control cows. Consequently, the treatment group for each feed additive consisted of 195 to 315 cows (15 cows for 13 to 21 farms, respectively). Additionally, an average of 370 cows per farm were randomly assigned to the control group (CG), resulting in CGs comprising 5550 to 7770 cows (for 13 to 21 farms, respectively). In summary, for each feed additive, there were a minimum of 195 cows in the treatment group and 5550 cows in the control group, in a fully multi-site randomized trial with unequal group sizes.

The data analysis followed two simulated scenarios: (1) a naive approach, which considers that feed additives are supplied indiscriminately to all farms without considering any efficacy predictions, and (2) an optimized approach, which assumes that feed additives are supplied only to farms with a high likelihood of being positively impacted, in terms of methane emissions (50% of the farms approximately), according to the AI-driven model. Thus, the data analysis is performed twice, the first time considering all the treatment groups (naive approach) for each feed additive, and the second time considering only farms with high response to the feed additives, in terms of methane emissions (optimized approach). Finally, the results of CG, NG, and OG groups are presented for each feed additive in the next tables.

Milk production data (liters/day) for 90 days following feed additive administration were obtained from the NOA system, which is used by approximately 95% of dairy farms in Israel. This automated system records and updates information on various parameters, including milk production (liters/day), protein (%), lactose (%), fat (%), and somatic cell count (thousand cells/ml). These parameters were used to analyze the effects of each feed additive.

### Animals and methane–mitigating feed additives

Israeli Holstein cows from 21 commercial dairy farms in Israel participated in this study. The herds were fed using a standard mixed diet (32/68 forage–to–concentrate ratio), and the cows of each herd were housed in the same barn. Farms participating in this study cover a diverse range of geographical settings within Israel, encompassing areas from arid deserts to cooler mountainous regions and more temperate plains, which helps mitigate the seasonal effects on milk production (Tsartsianidou et al., 2021).

In this study, we selected four feed additives based on essential oils: Agolin Ruminant (Altech, Biere, Switzerland), Allimax (Allimax Animal Health, Vaasen, Netherlands), RelyOn (Anavrin, Veto, Cadenazzo, Swiss), and Orego-Stim (Bar Magen, Ibadan, Nigeria), and one monensin–based feed additive, Kexxtone (Elanco, Indianapolis, USA). The doses used followed the manufacturer’s recommendations (1gram/cow/day for Agolin Ruminant and RelyOn, 2 grams/cow/day of Orego-Stim, 2.2 grams/cow/day of RelyOn, and 1 bolus/cow of Kexxtone). Since the farms analyzed are commercial, the local staff identified the treatment-group cows by their tags and supplied the methane–mitigating feed additives according to the manufacturer’s recommendations. No adverse events were reported.

### AI–driven model, ruminal samples, and sequencing

Briefly, the AI-driven model analyzes microbiome sequences to build farm–specific microbiome networks. It identifies and validates genetic markers associated with the targeted biological condition, such as methane emissions. Based on this analysis, the model generates an efficacy score that estimates the probability of a given feed additive positively influencing the targeted condition. Regarding to the microbiome sampling, the sample size question was dressed in a previous study (Altshuler et al., 2023; Altshuler et al., 2025), but a sample size of 15 cows could representatively predict the expected feed additive efficacy in terms of enteric methane emissions for a herd of up 3000 cows due to the absence of a significant correlation between farm size and the prediction engine’s accuracy.

The ruminal samples used by the model were obtained using a standard stomach tubing (polyvinyl chloride orogastric tube). Collected samples were sequenced using the shotgun sequencing method. Sequences were trimmed and cleaned producing sequences higher than 100 base pairs, with a Phred score higher than 35; then, the sequences were used to construct farm genetic networks (Altshuler et al., 2023).

### Milk production data and statistical analyses

Milk parameters were obtained from the Israeli Cattle Breeders Association (ICBA) through their dairy herd management program (NOA), which was designed to give the herd manager up- to-date information regarding all aspects of dairy activity, including herd management, feeding, and milk production recording and reports. All the data is registered in the national Herdbook database (ICBA, 2024). The NOA system was introduced in 2000 and is currently used in 95% of all dairy farms in Israel.

All statistical analyses were performed using PAST 4.17 software. Kruskal-Wallis test, with post hoc analysis using Mann-Whitney pairwise corrected by sequential Bonferroni, was used to compare treatments (control, naive, and optimized) according to each feed additive. The result tables present median values.

## RESULTS

Milk production in all the CGs was very similar, with a median fluctuating between 37.33 to 38.42 liters per day. In general, milk production significantly increased in each approach (naive and optimized). In the naive scenario (CG–NG *P* < 0.001, Table 1), milk production increased between 1% to 8.1%, with Orego–Stim producing the lowest increase (1%) being the only feed additive statistically like their control (*P* = 0.11), and RelyOn producing the highest increment of milk production (8.1%). Otherwise in the optimized scenario, the increase in milk production was even more remarkable (CG-OG *P* < 0.001, Table 1), fluctuating between 10% to 14.5%; thus, under the optimized approach, milk production reached values equal or higher than 42 liters per day with each feed additive. Interestingly, and contrary to the naive approach results, RelyOn, together with Agolin, produced the lowest increment and Orego–Stim producing the highest increment in milk production. In the case of RelyOn, the observed increase in milk production was very similar in both approaches (41.27 and 42 liters/day for naive and optimized approaches, respectively), being significantly higher than that obtained in the CG (38.19 liters/day). Figure 1 presents the average milk production for each feed additive across the experimental groups (CG, NG, and OG). Supplemental Material 1 details the data dispersion.

**Table 1.** Milk production (liters/day) according to feed additive and treatment.

| Feed Additive | Treatment<sup>1</sup> |  |  | *P* value<sup>2</sup> |  |  |
| --- | --- | --- | --- | --- | --- | --- |
|  | CG | NG<br>(change in %)<sup>3</sup> | OG<br>(change in %) | CG-NG | CG-OG | NG-OG |
| Agolin<br>(*n* = 16 sites)<sup>4</sup> | 38.42 | 39.29 (+2.2) | 42.27 (+10) | 0.001 | 1.37E-10 | 0.02 |
| Allimax<br>(*n* = 14 sites) | 37.59 | 39.03 (+3.8) | 42.26 (+12.4) | 0.004 | 5.55E-11 | 0.0003 |
| Kexxtone<br>(*n* = 21 sites) | 37.84 | 39.87 (+5.3) | 42.45 (+12.2) | 2.41E-07 | 1.08E-20 | 6.23E-06 |
| RelyOn<br>(*n* = 13 sites) | 38.19 | 41.27 (+8.1) | 42.00 (+10) | 1.57E-08 | 3.65E-07 | 0.527 |
| Orego-Stim<br>(*n* = 13 sites) | 37.33 | 37.70 (+1) | 42.76 (+14.5) | 0.11 | 9.43E-28 | 1.31E-9 |

<sup>1</sup>CG = control group; NG = naive group; OG = optimized group.
<sup>2</sup>Post-Hoc analysis with Mann-Whitney pairwise test, corrected by sequential Bonferroni.
<sup>3</sup> Percent of the change to CT; symbols indicate, + = increase; – = decrease.
<sup>4</sup> Number of farms tested for each feed additive.

**Figure 1.**
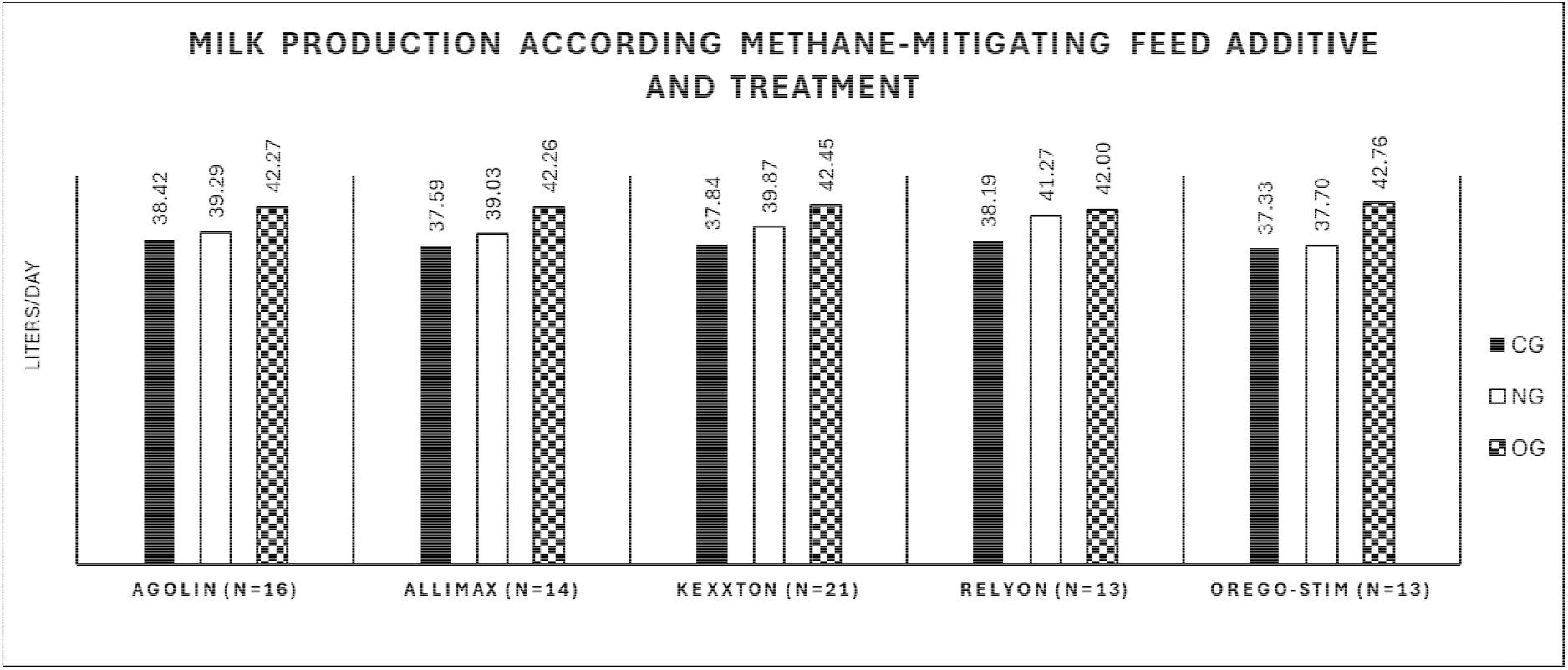
Milk production on average, according to feed additive and treatment. CG indicates the control group, NG: indicates the naive group, and OG: indicates the optimized group. The average value is at the top of each column.

Milk fat content in cows receiving the feed additives tended to be lower than their controls in all cases; however, in the naive approach, the fat reductions were between 1% to 3%, with only Kexxtone and RelyOn exhibiting significant differences compared to the CG (CG–NG, *P* = 0.0007 and P = 0.018 respectively, Table 2). In the case of the optimized approach, the reductions in fat content were more important, fluctuating in reductions of 1% to 6.2%, with Kexxtone producing the more pronounced decrease and Allimax presenting the lowest reductions, being in this case a non-significant reduction regarding to the CG (CG-OG *P* = 0.56).

**Table 2.** Fat content (%) according to feed additive and treatment.

| Feed Additive | Treatment<sup>1</sup> |  |  | *P* value<sup>2</sup> |  |  |
| --- | --- | --- | --- | --- | --- | --- |
|  | CG | NG<br>(change in %)<sup>3</sup> | OG<br>(change in %) | CG-NG | CG-OG | NG-OG |
| Agolin<br>(*n* = 16 sites)<sup>4</sup> | 3.86 | 3.82 (-1) | 3.72 (-3.6) | 1.0 | 0.0007 | 0.02 |
| Allimax<br>(*n* = 14 sites) | 3.97 | 3.91 (-1.6) | 3.93 (-1) | 0.35 | 0.16 | 0.56 |
| Kexxtone<br>(*n* = 21 sites) | 3.85 | 3.80 (-1.3) | 3.61 (- 6.2) | 0.0007 | 2.11E-115 | 0.0006 |
| RelyOn<br>(*n* = 13 sites) | 4.05 | 3.93 (-3) | 3.93 (-3) | 0.018 | 0.015 | 0.56 |
| Orego-Stim<br>(*n* = 13 sites) | 4.05 | 4.00 (-1.2) | 4.00 (-1.2) | 0.20 | 0.01 | 0.29 |

<sup>1</sup>CG = control group; NG = naive group; OG = optimized group.
<sup>2</sup>Post-Hoc analysis with Mann-Whitney pairwise test, corrected by sequential Bonferroni.
<sup>3</sup> Percent of the change to CT; symbols indicate, + = increase; – = decrease.
<sup>4</sup> Number of farms tested for each feed additive.

In terms of protein content, only minor variations were observed compared to the CG groups, with some cases showing reductions, while others exhibited an increase in protein content (Table 3). In the naive approach, cows receiving Agolin, Allimax, and Orego-Stim exhibited an increase in protein content (0.9%, 2%, and 0.6% respectively) regarding to the CGs, while cows receiving Kexxtone and Relyon decreased the protein content by 1.5% and 0.3%, respectively. Among the changes observed, only Allimax and Kexxtone were significantly different from their CGs (CG– NG; *P* = 0.002, and P = 0.0007 respectively). In the optimized approach, a general tendency for decreased protein content compared to the CG was observed; with Allimax being the only feed additive exhibiting a slight (0.9%), but still non-significant (*P* = 0.55) increase in protein content regarding the CG.

**Table 3.**
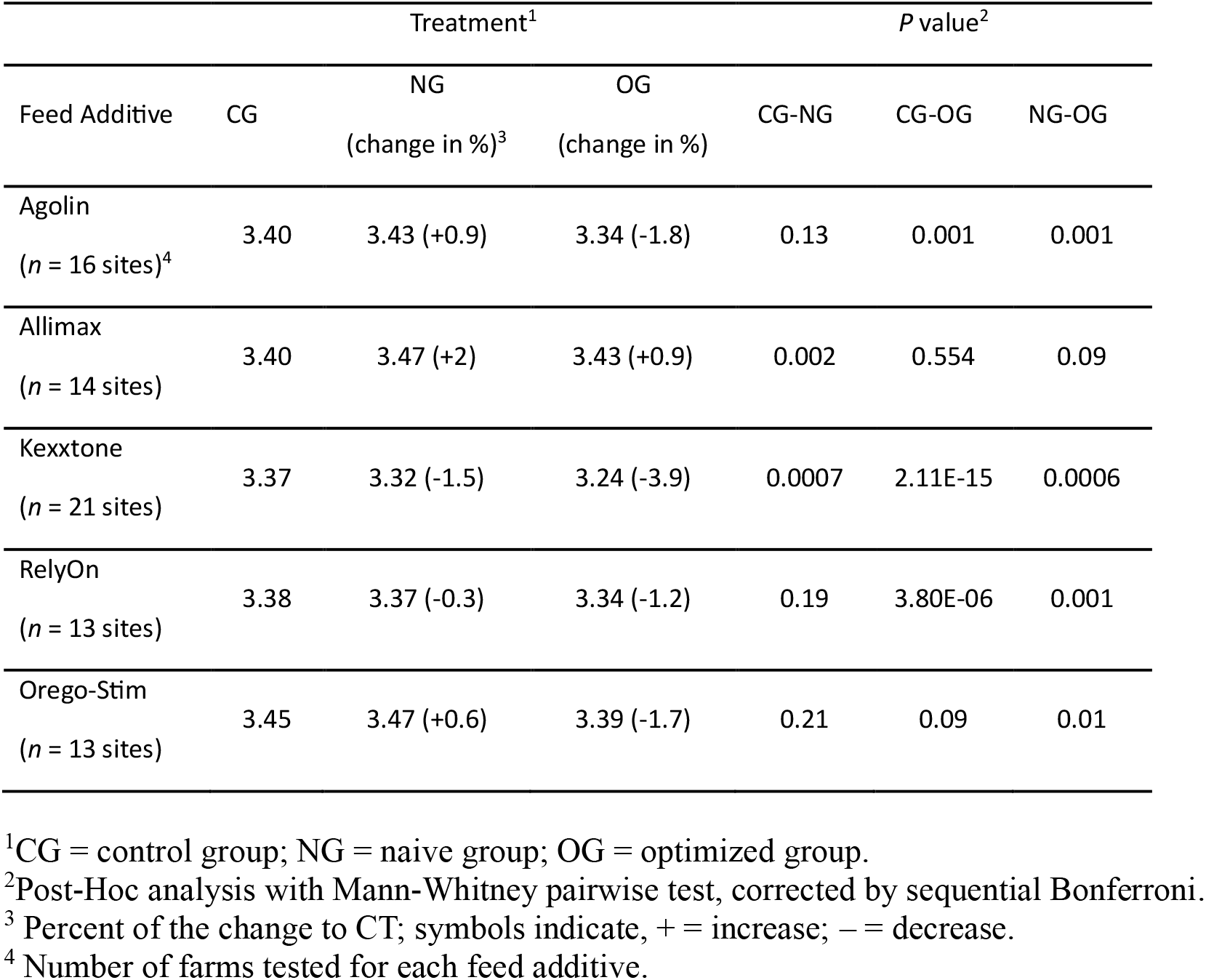
Protein content (%) according to feed additive and treatment.

The tested methane–mitigating feed additives minimally affected lactose content (Table 4). In both approaches (naive and optimized), no clear trends or significant changes were observed for most additives (*P* > 0.5), except for Allimax. Interestingly, Allimax significantly increased lactose content in both approaches, 0.8% (*P* = 1.67E-5) in the NG and 1.7% (*P* = 3.72E-6) in the OG group. Also, RelyOn exhibited an increase in lactose content in both naive and optimized approaches (0.4% and 0.2% respectively), but these increases were non-significant (*P* = 0.48 and *P* = 0.97 for each approach). Contrary, Agolin and Kexxtone exhibited a slight decrease in lactose content in both approaches, but again, this change respect to the CGs was non-significant (*P* = 0.67 and *P* = 0.15 respectively for the naive approach and *P* = 0.11 and *P* = 0.02 respectively for the optimized approach).

**Table 4.** Lactose content (%) according to feed additive and treatment.

| Feed Additive | Treatment<sup>1</sup> |  |  | *P* value<sup>2</sup> |  |  |
| --- | --- | --- | --- | --- | --- | --- |
|  | CG | NG<br>(change in %)<sup>3</sup> | OG<br>(change in %) | CG-NG | CG-OG | NG-OG |
| Agolin<br>(*n* = 16 sites)<sup>4</sup> | 4.78 | 4.77 (-0.2) | 4.76 (-0.4) | 0.67 | 0.11 | 0.28 |
| Allimax<br>(*n* = 14 sites) | 4.79 | 4.83 (+0.8) | 4.87 (+1.7) | 1.67E-5 | 3.72E-6 | 0.12 |
| Kexxtone<br>(*n* = 21 sites) | 4.79 | 4.77 (-0.4) | 4.77 (-0.4) | 0.15 | 0.02 | 0.37 |
| RelyOn<br>(*n* = 13 sites) | 4.77 | 4.79 (+0.4) | 4.78 (+0.2) | 0.48 | 0.97 | 0.60 |
| Orego-Stim<br>(*n* = 13 sites) | 4.84 | 4.85 (+0.2) | 4.83 (-0.2) | 0.35 | 0.17 | 0.64 |

<sup>1</sup>CG = control group; NG = naive group; OG = optimized group.
<sup>2</sup>Post-Hoc analysis with Mann-Whitney pairwise test, corrected by sequential Bonferroni.
<sup>3</sup> Percent of the change to CT; symbols indicate, + = increase; – = decrease.
<sup>4</sup> Number of farms tested for each feed additive.

Finally, the somatic cell content was evaluated. In both the naive and optimized approaches, a decrease in somatic cells was observed (Table 5). The decrease fluctuated between 12.8% and 40.4% for the naive and 7.5% and 34% for the optimized approach. Cows receiving Orego–Stim exhibited the highest decrease in somatic cell content for each approach, producing significant differences compared to the CG (*P* = 2.84E-7 and *P* = 1.93E-6 for naive and optimized approaches, respectively). In the case of Agolin, Kexxtone and RelyOn, the decrease in somatic cells content regarding CGs was non-significant in both approaches. Finally, Allimax exhibited a significant decrease in the naive approach (*P* = 0.006) but this decrease was non-significant in the optimized approach (*P* = 0.12).

**Table 5.** Somatic cell (thousand cells/ml) content according to feed additive and treatment.

| Feed Additive | Treatment<sup>1</sup> |  |  | *P* value<sup>2</sup> |  |  |
| --- | --- | --- | --- | --- | --- | --- |
|  | CG | NG<br>(change in %)<sup>3</sup> | OG<br>(change in %) | CG-NG | CG-OG | NG-OG |
| Agolin<br>(*n* = 16 sites)<sup>4</sup> | 219.58 | 170.47 (-22.4) | 187.92 (-14.4) | 0.012 | 0.82 | 0.33 |
| Allimax<br>(*n* = 14 sites) | 221.69 | 161.75 (-27) | 180.90 (-18.4) | 0.006 | 0.12 | 0.95 |
| Kexxtone<br>(*n* = 21 sites) | 228.98 | 189.44 (-17.3) | 211.75 (-7.5) | 0.15 | 0.80 | 0.73 |
| RelyOn<br>(*n* = 13 sites) | 226.28 | 197.30 (-12.8) | 184.46 (-18.5) | 0.21 | 0.34 | 0.58 |
| Orego-Stim<br>(*n* = 13 sites) | 242.53 | 144.46 (-40.4) | 160.07 (-34) | 2.84E-7 | 1.93E-6 | 0.25 |

<sup>1</sup>CG = control group; NG = naive group; OG = optimized group.
<sup>2</sup>Post-Hoc analysis with Mann-Whitney pairwise test, corrected by sequential Bonferroni.
<sup>3</sup> Percent of the change to CT; symbols indicate, + = increase; – = decrease.
<sup>4</sup> Number of farms tested for each feed additive.

## DISCUSSION

Overall, all tested feed additives led to an increase in milk production. This increase was significant in both treatment approaches, with a notably greater impact observed in the optimized approach. This increase was 5.3% and 11.2% on average for the naive and the optimized approaches, respectively. The results observed in this study, especially the results obtained with the NG, are in line with some meta-analyses reporting the positive effect of essential oils on milk production on different ruminants (Daning et al., 2021; Dorantes–Iturbide et al., 2022; Tudor et al., 2023). Similar to the essential oils, the tested monensin product also increased milk production, with the increase being consistent with previous reports (Becket et al., 1998; Ahvanooei et al., 2023).

The increase in milk production was generally accompanied by a tendency toward reduced fat content, which was statistically significant for most feed additives in the OG. Interestingly, Allimax showed only a slight, non-significant reduction in fat content compared to CG. In percentage terms, the reduction in fat content was approximately 1.5% in the naive approach and 3.9% in the optimized approach compared to CG. Other studies have reported diverse changes in fat content after the administration of essential oils or monensin products, with a slight increase in some cases (Ahvanooei et al., 2023; Becket et al., 1998; Daning et al., 2021), and no change in others (Benchaar et al., 2006; Tager and Kraus, 2011). The diversity of responses in previous studies could be based on the composition of each essential oil or monensin-tested product, the tested doses, the animal’s development stages, and the tested experimental conditions.

Regarding protein content, the observed responses can be divided according to the approach used in the analysis; thus, in the naive approach, no clear trends can be observed compared to CG. However, in the optimized approach, a trend of protein reduction is observed, suggesting that the protein synthesized may not increase proportionally to the increase in milk volume. Lactose content does not show significant changes regarding control; Only minor changes were observed between the naive and optimized approaches, but no consistent trend emerged. Finally, the somatic cell content decrease in all the tested additives and in each scenario. Interestingly, this product includes among its ingredients the active substance allicin, which is extracted from garlic (*Allium sativum*) and is known for its anti-inflammatory, antioxidant, antimicrobial, and antiparasitic activity (Ankri and Mirelman, 2016; Che et al., 2023), which could explain the similarities between NG and OG compared to CG.

The findings highlight the multifaceted benefits of methane-mitigating feed additives. Primarily designed to reduce methane emissions, these additives also demonstrate the potential to enhance milk production. This improvement can be further optimized using tools like our predictive AI-driven model, fostering a precision agriculture approach. Numerous studies have emphasized the necessity of integrating advanced technologies in the dairy industry, as they are essential for improving health and efficiency in milk production, supporting the development of sustainable, animal-friendly, and efficient livestock systems (Bianchi et al., 2022; de Vries et al., 2023; de Oliveira et al., 2024). Despite some observed drawbacks, such as reduced fat content, a preliminary analysis (Supplemental Material 2) shows that feed additives remain highly profitable, which added to the above, contribute to a more informed decision-making process.

Despite the extensive research on the effects of feed additives, most studies are conducted under highly controlled conditions or involve a limited number of animals. This underscores the need for large-scale investigations carried out under typical farm conditions to comprehensively assess the diverse impacts of feed additives on milk and meat production. Addressing this gap, the present study represents the first attempt to evaluate multiple methane-mitigating feed additives in situ across a substantial number of commercial dairy farms. The participating farms accurately mirror real-world conditions, ensuring the findings are both broadly representative and highly reliable.

## CONCLUSIONS

This study showed an increase in milk yield when different methane–mitigating feed additives were supplied to experimental groups in commercial dairy farms. The increase in milk production was higher in the optimized scenario, where the AI–driven model predicted a higher impact of the feed additives. Some milk parameters, such as fat content, were slightly affected when compared to CG; however, the results suggest that the positive effects of the feed additives were higher than the negative ones. The scope of this study, in terms of the animals tested and the number of participating farms, the number of methane–mitigating feed additives tested, and the data collection from an automated system, guarantees high confidence in the results and conclusions. The integration of the AI–driven model proved beneficial in helping farmers makes well-informed decisions.

## Supplemental Material

Supplemental material is available at https://doi.org/10.5281/zenodo.14197687.

## Notes

This study received no external funding, and the authors have not stated any conflicts of interest. Ethical approval was not necessary for this study. We sincerely thank the farmers from the participating dairy farms in Israel for their cooperation and support throughout this study. Their involvement and support were essential to the success of this research.

## Nonstandard abbreviations used

CG: control group
NG: naive group
OG: optimized group
NOA: dairy herd management program of the Israeli Cattle Breeders Association.

