## Supplementary Material 1 for "ANALYSIS OF THE IMPACT OF FIVE METHANE–MITIGATING FEED ADDITIVES ON MILK PRODUCTION AND ASSOCIATED PARAMETERS ACROSS MULTIPLE COMMERCIAL DAIRY FARMS"

**Supplementary Material 1.** Box plots of milk production according to feed additive and approach (naive and optimized). Y- axis expressed in liters/day

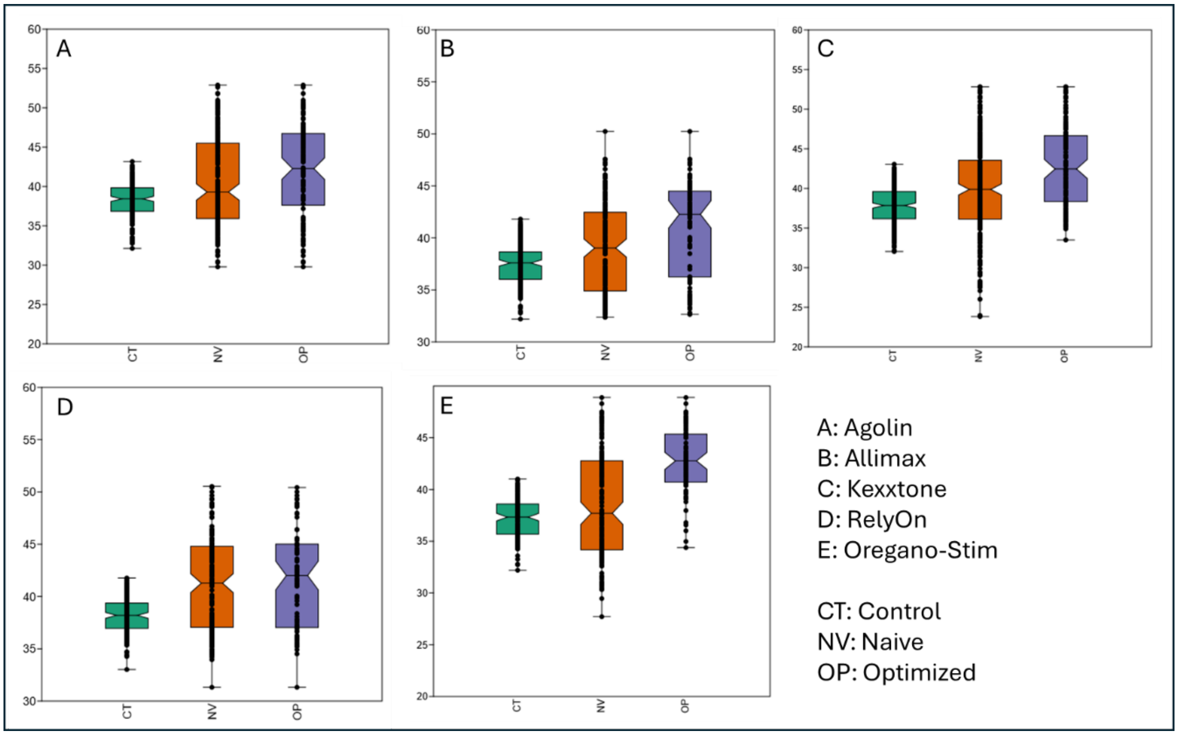
