## Supplementary Material 2 for "ANALYSIS OF THE IMPACT OF FIVE METHANE–MITIGATING FEED ADDITIVES ON MILK PRODUCTION AND ASSOCIATED PARAMETERS ACROSS MULTIPLE COMMERCIAL DAIRY FARMS"

**Supplementary Material 2.** Average energy corrected milk (ECM) according to feed additive and annual ECM (305 days of lactation period). The change of each additive to their control is indicated in brackets and expressed in dollars assuming USD 0.44/litter. CG indicates the control group; NG indicates the naive group, and OG indicates the optimized group.

| **Feed Additive** | **ECM** | | | **Annual ECM. Change to CT in brackets and expressed in dollars** | | |
|---|---|---|---|---|---|---|
| | **CG** | **NG** | **OG** | **CG** | **NG** | **OG** |
| **Agolin** (n = 16 sites) | 41.2 | 43.4 | 44.3 | 12561 | 13230 ($294) | 13499 ($412) |
| **Allimax** (n = 14 sites) | 40.6 | 42.1 | 44.0 | 12382 | 12833 ($198) | 13406 ($451) |
| **Kexxtone** (n = 21 sites) | 40.5 | 42.3 | 44.3 | 12362 | 12913 ($242) | 13520 ($509) |
| **RelyOn** (n = 13 sites) | 42.0 | 44.7 | 44.8 | 12818 | 13647 ($365) | 13675 ($377) |
| **Orego-Stim** (n = 13 sites) | 40.9 | 42.1 | 46.1 | 12488 | 12826 ($149) | 14068 ($695) |
